# Identification and validation of a non-genetically encoded vulnerability to XPO1 inhibition in malignant rhabdoid tumors – expanding patient-driven discovery beyond the N-of-1

**DOI:** 10.1101/2021.10.02.462793

**Authors:** Lianna J. Marks, Daniel Diolaiti, Prabhjot Mundi, Ervin S. Gaviria, Allison R. Rainey, Darrell J. Yamashiro, Ladan Fazlollahi, Hajime Hosoi, Yoshiki Katsumi, Yasumichi Kuwahara, Filemon S. Dela Cruz, Andrea Califano, Andrew L. Kung

## Abstract

Malignant rhabdoid tumors (MRTs) are rare, aggressive pediatric solid tumors, characterized by a 22q11 deletion that inactivates the SMARCB1 gene. Outcomes remain poor despite multimodality treatment. MRTs are among the most genomically stable cancers and lack therapeutically targetable genetic mutations. We utilized metaVIPER, an extension of the Virtual Inference of Protein-activity by Enriched Regulon (VIPER) algorithm, to computationally infer activated druggable proteins in the tumor of an eight month old patient and then expanded the analysis to TCGA and TARGET cohorts. *In vitro* studies were performed on a panel of MRT and atypical teratoid/rhabdoid tumor cell lines. Two patient-derived xenograft (PDX) mouse models of MRT were used for *in vivo* efficacy studies. MetaVIPER analysis from the patient’s tumor identified significantly high inferred activity of nuclear export protein Exportin-1 (XPO1). Expanded metaVIPER analysis of TCGA and TARGET cohorts revealed consistent elevations in XPO1 inferred activity in MRTs compared to other cancer types. All MRT cell lines demonstrated baseline activation of XPO1. MRT cell lines demonstrated *in vitro* sensitivity to the XPO1 inhibitor, selinexor which led to cell cycle arrest and induction of apoptosis. Targeted inhibition of XPO1 in patient-derived xenograft models of MRT using selinexor resulted in abrogation of tumor growth. Selinexor demonstrates efficacy in preclinical models of MRT. These results support investigation of selinexor in a phase II study in children with MRT and illustrate the importance of an N-of-1 approach in driving discovery beyond the single patient.

**Statement of Translational Relevance:** We describe the patient-driven discovery of XPO1 activation as a non-genetically encoded vulnerability in MRTs. The application of metaVIPER analysis to tumors lacking actionable oncogenic alterations represents a novel approach for identifying potential therapeutic targets and biomarkers of response. Our preclinical validation of selinexor confirms XPO1 inhibition as a promising therapeutic strategy for the treatment of MRT.

## Introduction

Comprehensive genomic and transcriptomic profiling of tumors has the potential to impact patient care and guide treatment decisions for individual patients, especially those with rare cancers lacking efficacious conventional therapeutic options (1). Malignant rhabdoid tumors (MRTs) are rare, aggressive pediatric tumors associated with extremely poor prognosis. MRTs are characterized by the biallelic inactivation of the *SMARCB1* gene (also known as *SNF5, INI1*, or *BAF47*) through chromosomal deletions and nonsense, missense, or frameshift mutations, with as many as 35% of cases associated with germline variants in *SMARCB1* (2,3). MRTs occur throughout the body but are most frequently localized to the kidney and the brain, where they are referred to as atypical teratoid/rhabdoid tumors (ATRTs). The peak incidence of MRT is in infancy, with a median age of 11 months at diagnosis (4). Despite multimodal therapy, outcomes remain poor with an overall 4-year survival of 23.2%, and only 8.8% in patients younger than six months (5).

Compared to adult tumors, genomic characterization of pediatric tumors generally demonstrates few single nucleotide variants (SNV) and low overall tumor mutational burden (6). Furthermore, MRTs are one of the most genomically stable of all pediatric cancers with a mutation rate of only 0.084 per megabase, which is 10 to 50-fold lower than the majority of adult tumors (7). SMARCB1, characteristically inactivated in MRT, is a core subunit of the highly conserved nucleosome-remodeling SWItch/Sucrose Non-Fermentable (SWI/SNF) complex which regulates nucleosome positioning and occupancy in an ATP-dependent manner, thereby controlling DNA accessibility (8). Within the SWI/SNF complex, SMARCB1 is required for efficient binding to DNA, transcriptional activation of enhancers of lineage-specific genes, and resolution of bivalency at promoters of lineage-specific genes (9). Loss of SMARCB1 results in gene expression changes that drive tumorigenesis by multiple mechanisms, including epigenetic perturbations of differentiation and the cell cycle (9). Due to the lack of targetable genomic alterations in MRT, new methods for discovering candidate, non-genetically-encoded tumor vulnerabilities are needed.

Both genetic and epigenetic alterations, in combination with complex networks that regulate gene expression and post-translational modifications, contribute to dysregulated protein activity (10). The Virtual Inference of Protein-activity by Enriched Regulon analysis (VIPER) algorithm is a computational tool for the inference of protein activity in cancer at the individual tumor level (11,12). The ability of VIPER to accurately and reproducibly infer differential protein activity has been extensively validated in several contexts: 1) to identify candidate ‘Master Regulators’ (MRs) whose aberrant activity is critical to tumor viability and growth *in vivo*; 2) to identify regulatory proteins whose differential activity associates with tumor subtypes, metastatic state, treatment sensitivity, mechanisms of resistance, and prognosis; and 3) to identify drugs/combinations that disrupt the regulatory proteins responsible for a tumor’s homeostatic state as a predictor of drug sensitivity (13-19).

We report the case of an infant diagnosed with MRT, whose tumor was assessed using whole exome and RNA sequencing. Using the VIPER algorithm in a N-of-1 application, we identified aberrant XPO1 activity by comparison to other cancer types. Leveraging this patient-driven observation, we demonstrate XPO1 inhibition as a class-specific vulnerability for MRTs, suggesting a new therapeutic approach for this disease.

## Results

### Case Report

A previously healthy eight month old girl presented with a firm abdomen and was found to have a 10 cm right hepatic mass on ultrasound. An abdominal computed tomography (CT) scan showed a 10.1 × 7.2 × 11.5 cm lobulated heterogeneous mass arising from the right hepatic lobe (Figure 1A). She was also noted to have multiple small pulmonary nodules consistent with metastatic disease on chest CT. Laboratory workup demonstrated an alpha feto-protein (AFP) of 71.5 ng/ml (mean AFP for age 8ng/ml, normal range 0.8-87ng/ml)(20). Histopathological evaluation of an incisional biopsy of the liver mass showed small cell morphology (Fig. 1B) with immunohistochemistry (IHC) showing loss of INI1 protein expression (Fig. 1C) (21). In the absence of rhabdoid morphology the tumor was classified as small cell undifferentiated (SCUD) hepatoblastoma. Under the current pediatric liver tumor classification, in the absence of rhabdoid morphology INI1 negative IHC is insufficient to diagnose MRT, however, recent studies have shown that presence of INI1 negativity with IHC in a pediatric liver tumor is diagnostic of MRT even in the absence of rhabdoid morphology (22-24). Given these results, a presumptive diagnosis of high risk hepatoblastoma was rendered due to non-elevated AFP and metastatic disease status. The patient was enrolled on Children’s Oncology Group (COG) study AHEP0731, high-risk stratum, and received 2 cycles of vincristine, irinotecan, and temsirolimus (VIT) and 4 cycles of cisplatin, 5-fluorouracil, and vincristine (C5VD) prior to a right partial hepatectomy.

**Figure 1:**
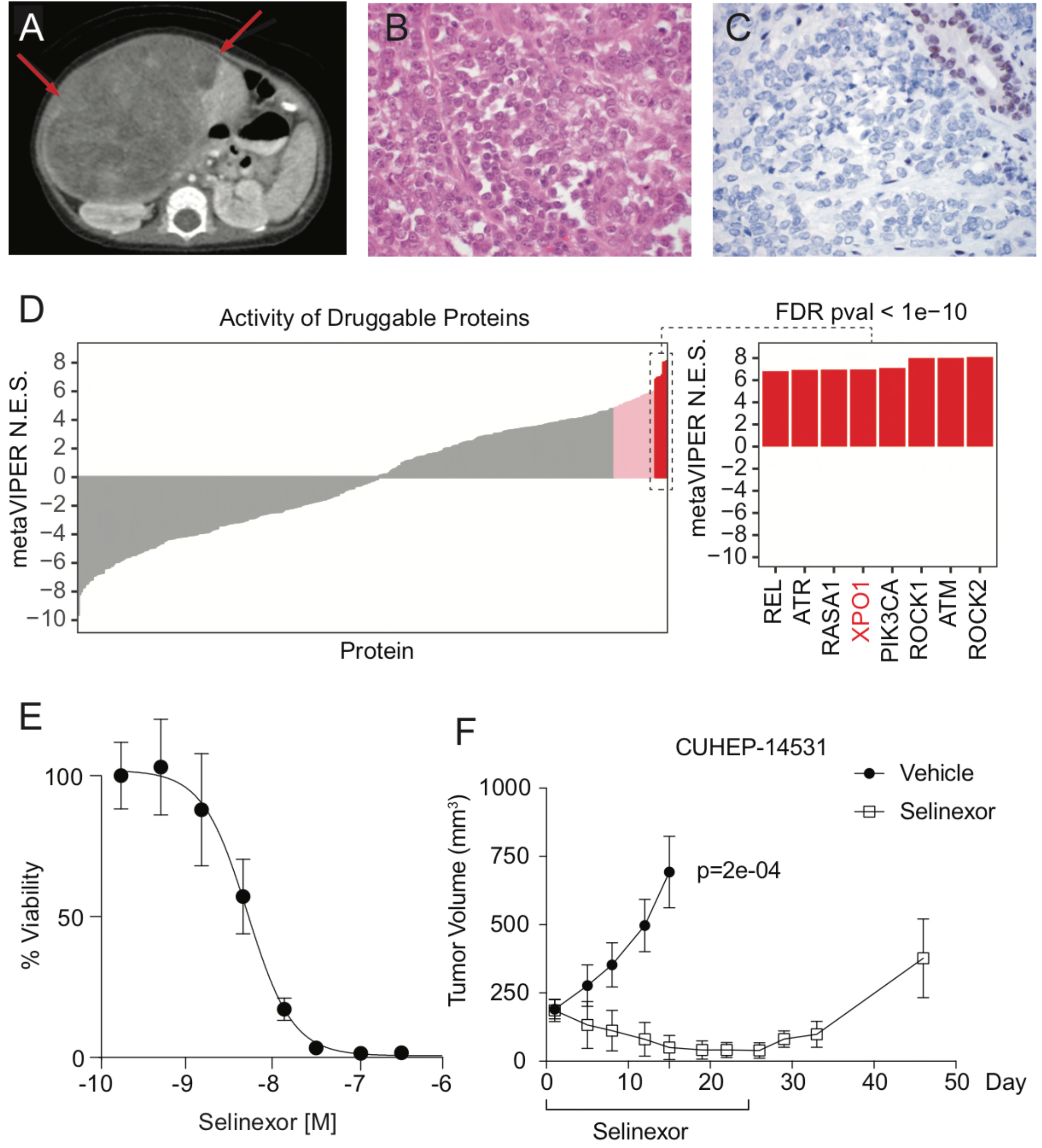
Clinical presentation, MetaVIPER analysis, and sensitivity to small molecule inhibitor selinexor in patient-derived pre-clinical model. **A)** Diagnostic imaging demonstrating the presence of a large hepatic tumor. **B)** Hepatic tumor biopsy with H&E stain demonstrates small cell morphology. 600x **C)** INI1 immunohistochemistry stain highlights an entrapped bile duct (positive internal control), tumor cells are negative. 600x **D)** MetaVIPER inferred activity of all ‘druggable’ proteins in the patient tumor biopsy from which the CUHEP-14531 PDX model was developed. The inset to the right highlights the top 8 most aberrantly activated targets. **E)** IC_50_ of MRT cells cultured from CUHEP-14531 PDX and treated with selinexor for 72 hours. **F)** Tumor response in CUHEP-14531 PDX (n=8 per arm) treated with selinexor.

Review of the surgical resection specimen demonstrated focal areas of large polygonal tumor cells with eccentric nuclei and dense eosinophilic cytoplasm consistent with a rhabdoid morphology. For further evaluation, whole exome sequencing (WES) and RNA sequencing were performed and copy number analysis confirmed homozygous deletion of *SMARCB1* on chromosome 22q11.23 with consequent complete loss of mRNA expression (25). Based on this information the tumor was reclassified as an MRT (21,23-25). WES did not identify any additional known oncogenic variants.

Following 6 total cycles of chemotherapy, the patient experienced on therapy progression in the lungs prompting removal from trial therapy. She subsequently received empiric ifosfamide and etoposide alternating with vincristine, doxorubicin, and cyclophosphamide for five cycles with minimal benefit, followed by metastatecomy of four lung lesions. This was followed by high dose chemotherapy with melphalan, carboplatin, etoposide, and autologous stem cell rescue, whole lung (1050 cGy) and abdomen (1200 cGy) radiation, and repeat gross total resection, but the patient ultimately succumbed to inexorable disease progression.

### Increased XPO1 Activity in Patient’s Tumor

Due to the lack of targetable genomic alterations from tumor WES profiling, we utilized the VIPER algorithm to infer protein activity from the transcriptome profile. VIPER leverages cancer type-specific gene regulatory networks, such as those constructed by ARACNe (Algorithm for the Reconstruction of Accurate Cellular Networks), with sample-specific gene expression signatures. It infers differential protein activity by using the expression of tens to hundreds of target genes of each regulatory protein as a robust and sensitive reporter for its activity (11,12,26). Because there was an insufficient number of MRTs profiled in the TARGET database to generate an MRT-specific network, we utilized metaVIPER (27), a recently published extension of VIPER that can effectively infer protein activity by integrating the results of VIPER using all available cancer networks at the protein level. metaVIPER analysis was used to computationally infer the differential activity of 412 proteins currently classified as ‘druggable’ using the patient’s gene expression profile. This approach identified several proteins with elevated inferred activity, including key regulators of actin cytoskeleton and cell polarity (ROCK1/2), DNA damage response (ATM and ATR), cell signaling (PIK3CA and BRAF), and protein trafficking (XPO1) (Fig. 1D and Supplementary Table 1). XPO1 is a nuclear export protein that selectively transports tumor suppressor and growth regulatory proteins, including protein 53 (P53), retinoblastoma 1 (RB1), and cyclin-dependent kinase inhibitor 1B (CDKN1B, p27^Kip1^), out of the nucleus and inhibits their function (24,25). An emerging class of small-molecule orally bioavailable XPO1 inhibitors, the selective inhibitors of nuclear export (SINE), have already shown preclinical efficacy in several hematologic and solid malignancies and recently gained FDA approval for the treatment of multiple myeloma (28). Selinexor is currently in early phase trials in pediatrics (29,30).

### Effects of Selinexor Treatment in Patient-Specific PDX Model

A patient-derived xenograft (PDX) model (CUHEP-14531) was generated using tumor material obtained from the patient’s biopsy specimen and confirmed to harbor the *SMARCB1* deletion (Supplementary Table 2). Cells harvested from the PDX model were expanded and treated with selinexor *ex vivo* under short-term culture conditions. Results from a dose-response assay revealed that cells were highly sensitive to selinexor (IC_50_ of 5.2 nM 95% CI 4.2-6.3nM) (Fig. 1E) in comparison to multiple myeloma cells (median IC_50_ 165 nM in 12 cell lines) for which selinexor has been granted FDA approval (31). Additionally, pharmacokinetic studies of selinexor have shown that the FDA-approved dose of 80 mg twice a week achieves a peak plasma concentration of 1-1.5 uM suggesting *in vitro* sensitivity at clinically achievable levels (29,30).

We next assessed the *in vivo* efficacy of selinexor in a PDX model derived from the case patient’s tumor (CUHEP-14531). Treatment of mice with established MRT tumors demonstrated tumor regression over the 25 days of treatment with selinexor, however, the abrogation of tumor growth was dependent upon sustained treatment (Fig. 1F). The magnitude of response to selinexor was comparable to standard of care therapy with ifosfamide and etoposide (Fig. S1). Unfortunately, the patient was unable to enroll in the selinexor pediatric trial ADVL1414 (NCT02323880) due to body surface area requirements and lack of oral suspension.

### Increased XPO1 Activity in MRT

To evaluate the relevance of XPO1 as a target across MRTs, we performed metaVIPER analysis on TCGA and selected TARGET tumor types (Fig 2A). Notably, XPO1 was a significantly activated protein in 94% of MRTs (64 of 68 samples with FDR p-value < 0.01). Based on both within sample protein normalized enrichment score (NES) (Fig 2A) and rank (Fig. S2A), the overall XPO1 inferred activity in MRTs was higher compared to most other cancers. Other tumors with high XPO1 inferred activity included Wilms tumor, testicular germ cell tumors, esophageal carcinoma, and acute myeloid leukemia (AML) (Figs. 2A, S2A). The median rank of XPO1 activity in MRTs was 6.5, making it the third most frequently activated druggable protein across MRTs, behind DNA topoisomerase II alpha (TOP2A) and dual specificity protein kinase TTK, a protein associated with cell proliferation and chromosome alignment (Fig. S2B). Additional recurrently activated therapeutic targets identified in MRTs include the purine biosynthesis enzyme GART, the kinesin-like spindle protein KIF11, Haspin (GSG2), and DNMT3B (Fig. S2B). Unlike most other tumor types, in which XPO1 activity was distributed over a wide range of values, MRTs have a relatively high and uniform XPO1 inferred activity (Fig. 2A: IQR for NES 4.59 to 6.08).

**Figure 2:**
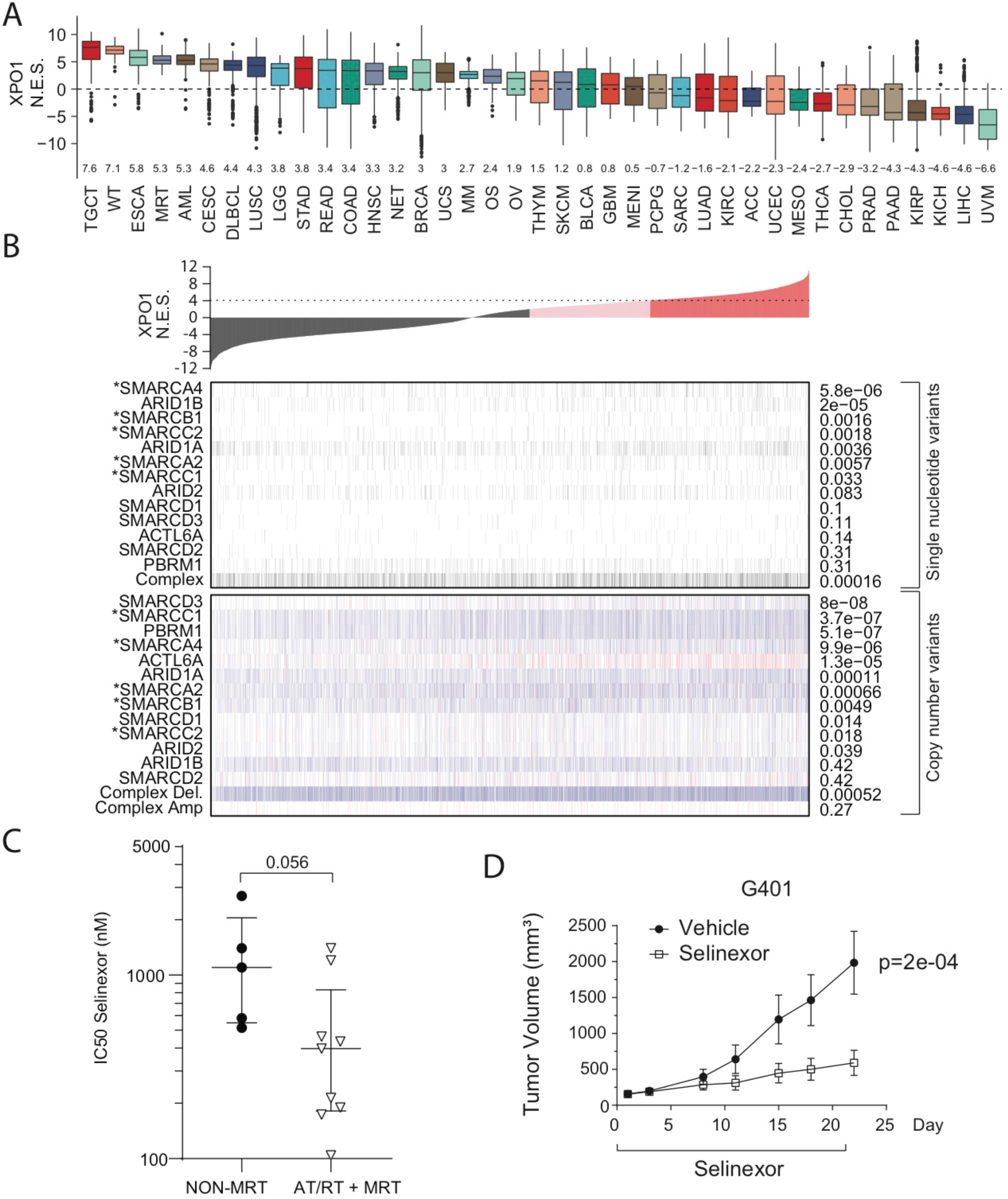
MetaVIPER inference of XPO1 activity and sensitivity of rhabdoid cell lines to the XPO1 small molecule inhibitor selinexor. **A)** Boxplots representing the distribution of metaVIPER inferred XPO1 activity for 33 TCGA tumor cohorts, 3 TARGET pediatric tumor cohorts (MRT, Wilms, osteosarcoma), and a neuroendocrine tumor cohort. The median and interquartile range for NES values is represented by each box for the respective tumor cohort. NES values from enrichment analysis are comparable to Z-scores, with higher scores representing increased activity. **B)** Co-segregation of SNV and CNV events in SWI/SNF complex genes with XPO1 activity. Top: Barplot of XPO1 activity in each TCGA tumor. Middle: SNVs in SWI/SNF complex genes. SNVs in core subunits, such as SMARCA4 and SMARCB1, and ARID1B most strongly co-segregate with XPO1 activation. Estimated p-values from enrichment analysis are to the right. Bottom: CNVs are far more common than SNVs in SWI/SNF complex genes. Heterozygous (light blue) and homozygous (dark blue) deletions in several SWI/SNF complex genes co-segregate with increased XPO1 activity, as do amplifications (red) in ACTL6A. However, SWI/SNF deletions do not universally correspond with high XPO1 activity, as can be seen in several tumors towards the left of this plot. **C)** Dot plot showing IC_50_ of selinexor in MRT and ATRT vs non-MRT cell lines following 72 hours of treatment. **D)** Tumor response in G401 mouse xenograft (n=8 per arm) treated with selinexor. TCGA – the Cancer Genome Atlas; TARGET – Therapeutically Applicable Research to Generate Effective Treatments; NES – normalized enrichment score. MRT – malignant rhabdoid tumor; ATRT – atypical teratoid/rhabdoid tumor.

To gain insight into whether genetic disruption of the SWI/SNF complex is associated with aberrant XPO1 activity, we performed co-segregation analysis between genomic events, i.e., somatic single nucleotide variants (SNVs) and copy number variants (CNVs), and XPO1 activity using 33 cancer cohorts in TCGA (N=8,348 tumors). Across this pan-cancer atlas, XPO1 activity was found to co-segregate with SNVs (p=0.00016) and deletions (p=0.00052) in genes encoding the SWI/SNF complex subunits (Fig. 2B). Notably, XPO1 activation occurs in association with mutations in SWI/SNF complex “core” subunits, including SMARCB1, plus the AT-rich DNA binding subunits ARID1A and B, but not with accessory subunits (Fig. 2B) (8), suggesting that a broader correlation exists between loss of SWI/SNF function and increased XPO1 activity.

### MRT/ATRT cell lines demonstrate XPO1 activation and differential sensitivity to selinexor

To experimentally validate the *in silico* results above, we assembled a panel of MRT and AT/RT cell lines and utilized metaVIPER analysis to compare their XPO1 inferred activity to reference cell lines profiled in the Cancer Cell Line Encyclopedia (CCLE). Our analysis confirmed high XPO1 activity (Fig. S2C) in the selected cell lines, indicating that MRT/ATRT cell lines recapitulated the XPO1 activation observed in patient tumors. Protein expression of XPO1 was variable across a range of MRT/ATRT and other cancer cell lines and had overall higher expression compared to selected non-cancer cell lines (Fig. S2D). As expected, all MRT/ATRT cell lines were SMARCB1 negative, while the non-MRTs expressed SMARCB1 (Fig. S2D).

*In vitro* sensitivity to selinexor was then evaluated across the same panel of MRT/ATRT cell lines. The median IC_50_ for MRT cell lines following 72 hour treatment with selinexor was 200 nM (IQR 170-440 nM) for MRTs, 460 nM (IQR 400 nM-1.4 uM) for ATRTs, and 1.1 uM (IQR 580 nM-1.4 uM) for 5 non-MRT cell lines (Fig. 2C), demonstrating differential sensitivity of MRT cell lines to XPO1 inhibition compared to non-MRT cell lines. Next, we generated a tumor xenograft MRT model utilizing the G401 established MRT cell line. Treatment with selinexor over a period of 21 days significantly inhibited tumor growth (TGI= 71%) (p=0.0002) (Fig. 2D), confirming the sensitivity of MRT cells to selinexor *in vivo*.

### Selinexor inhibits XPO1 activity and induces cell cycle arrest and apoptosis

To gain insight on the mechanism of action of selinexor we treated selected MRT cell lines with vehicle (DMSO) or with a sublethal concentration of selinexor (30 nM) and performed post-perturbation RNAseq profiling and metaVIPER analysis to evaluate differential protein activity induced by the drug. This analysis demonstrated a predictable and significant decrease in XPO1 activity that became more profound between 6 and 24 hours in all four cell lines, and, in turn, a compensatory increase in XPO1 mRNA abundance (Fig 3A). In addition to confirming that the analysis is sufficiently sensitive to infer changes in XPO1 activity as a consequence of pharmacologic inhibition, it also highlights other potential mechanistic targets of selinexor (Fig. 3A, Fig. S3A). Next, we used the differential protein activity signature computed by metaVIPER and ran pathway enrichment analysis to determine which cancer hallmark pathways were most significantly impacted at this sublethal concentration of selinexor. We observed a marked downregulation in pathways involving cell cycle progression (E2F targets, G2M checkpoint), downregulation in the MYC pathway, and an increase in epithelial-mesenchymal transition, TNF-alpha signaling, and KRAS signaling (Fig. 3B).

**Figure 3:**
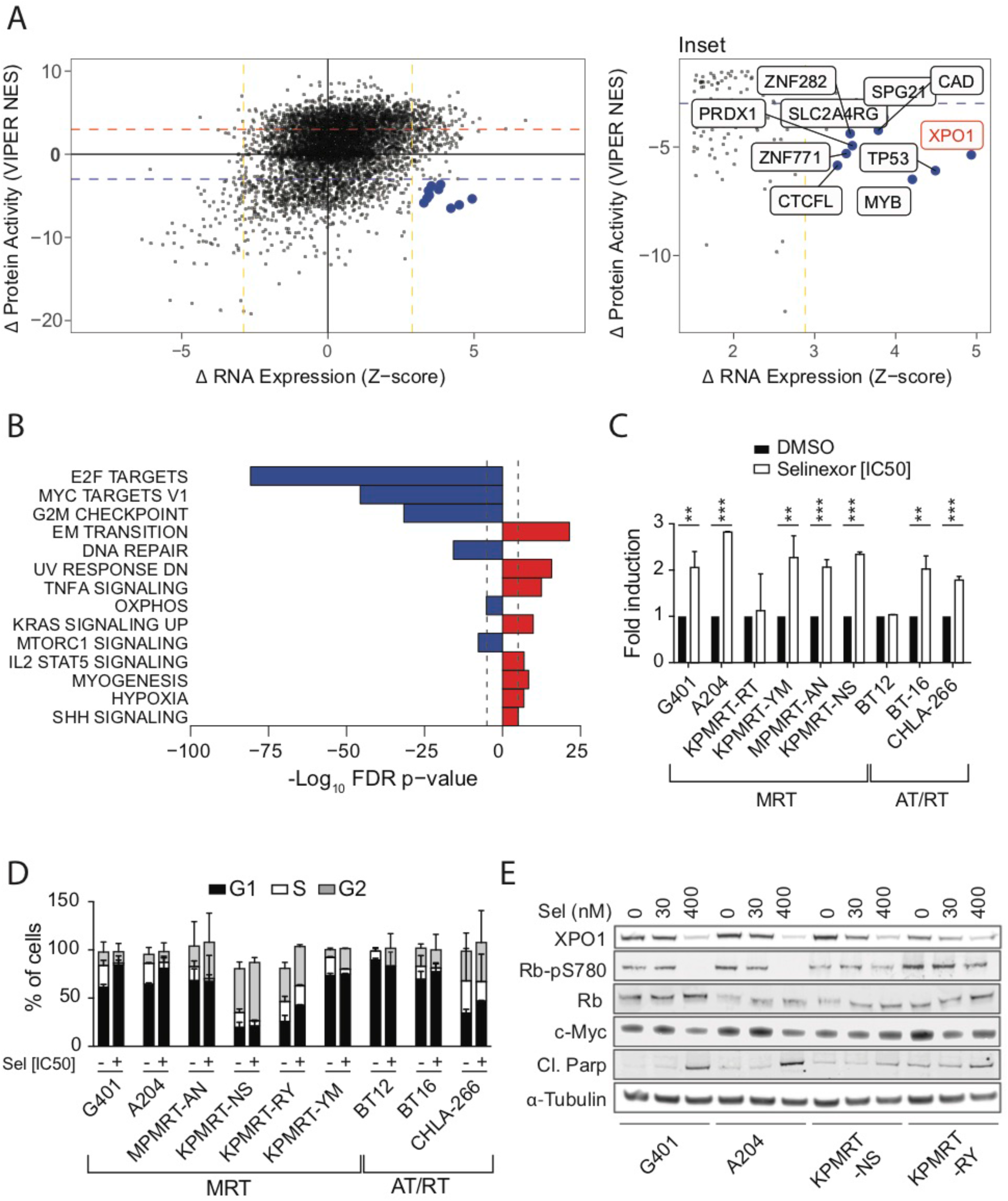
Pharmacodynamic analysis of selinexor in rhabdoid tumor cell lines using next generation sequencing, flow cytometry, and western blot. **A)** RNA-Seq was performed on four rhabdoid cell lines at baseline and after treatment with selinexor 30 nM at 6 and 24 hours. Changes in mRNA abundance positively correlate with MetaVIPER inferred activity at 24 hours, but XPO1 is amongst a small group of proteins (inset) whose activity is markedly decreased by selinexor in spite of significant compensatory increase in mRNA. **B)** The differential protein activity signature induced by selinexor—i.e., the inferred change in protein activity of about 6000 proteins including XPO1, was further analyzed using pathway enrichment analysis on the cancer hallmarks set from MSigDB. The pathways that are most significantly deactivated are in blue, while pathways with significantly increased activity after selinexor perturbation are in red. **C)** Apoptosis assay of cell lines treated with DMSO or 200 nM selinexor for 48 hours. **D)** Cell cycle analysis across MRT/ATRT cell lines treated with DMSO or 400 nM selinexor for 48 hours showing cell cycle arrest with a decreased percentage of cells in S phase and increased in G1 and G2/M. **E)** Immunoblot showing expression levels of the indicated proteins in G401, A204, KPMRT-NS and KPMRT-RY cell lines treated with selinexor at varying concentrations for 48 hours. Mean and standard deviation of three independent experiments is shown in (C) and (D). *=p<0.05, **=p<0.01, ***=p<0.001

In addition, we examined changes in the inferred activity of other druggable proteins whose activity strongly co-segregated with XPO1 in MRTs, including several epigenetic regulators such as DNMT1, EZH2, KDM1A, EP300, and BRD4, as well as several proteins involved in DNA repair (Fig. S3B). Interestingly, inhibiting XPO1 led to a loss of activity in the majority of these proteins suggesting a functional interdependency. Alternatively, there was increased activity of BRAF, the Rho-associated kinases ROCK1/ROCK2, and cereblon (CRBN), the target of lenalidomide in the E3-ubiquitin ligase pathway (Fig. S3B).

Consistent with the observed gene expression/protein activity changes, selinexor treatment of MRT/ATRT cell lines *in vitro* demonstrated induction of apoptosis at various levels in most cells tested (Fig. 3C) and cell cycle arrest with a decrease in the percentage of cells in S phase and reciprocal increase in the percentage of cells in G1 and G2/M phase (Fig. 3D). We then evaluated the protein levels and the cellular localization of XPO1 and XPO1 target proteins mediating cell cycle and apoptosis. Analysis of protein expression following selinexor treatment in 4 MRT cell lines showed decreased XPO1 expression (Fig. 3E). We also observed decreases in phosphorylation of RB (Fig. 3E), confirming that selinexor alters the expression of major cell cycle regulators. In parallel, we observed increased levels of cleaved PARP, consistent with an induction of apoptosis in MRT cells upon treatment with selinexor (Fig. 3E). Subcellular localization of XPO1 targets following selinexor treatment showed increased nuclear sequestration of mediators of cell cycle arrest p53, p27, and p21 (Fig. S3C), confirming that selinexor blocked the ability of XPO1 to shuttle its targets across the nuclear membrane.

### Effects of Selinexor Treatment in PDX MRT Model

Although cell lines were used to evaluate selinexor sensitivity of MRT *in vitro*, the mutational analysis performed using MSK-IMPACT (32), our institutional next-generation sequencing targeted panel, showed that most cell lines, including G401, harbored mutations in *TP53*, which are not common in rhabdoid patient tumors (Supplemental Table 2). Therefore, we assessed the *in vivo* effects of selinexor in a second MRT PDX model. Model MSKMRT-31222 was generated from primary tumor tissue obtained from the resection of a pelvic soft tissue mass in a 16 year old girl with treatment-refractory MRT. The model had absence of INI1 staining by IHC, and genomic characterization confirmed homozygous deletion of 22q11.23 involving *SMARCB1* with resultant loss of mRNA expression, but no mutations in *TP53* (Supplementary Table 2).

Treatment of MSKMRT-31222 PDX tumors with selinexor demonstrated an anti-tumor effect with stabilization of tumor volume (Fig. 4A). Similar to *in vitro* treatment of MRT cell lines, analysis of CUHEP-14531 PDX tumors collected after one week of selinexor treatment showed decreased XPO1, RB1-pSer780, and c-MYC (Fig. 4B) consistent with the anti-proliferative response observed *in vitro*. MSKMRT-31222 PDX tumors showed only a significant decrease in c-MYC (Fig. 4C). Interestingly, while the CUHEP-14531 tumors also demonstrated increased p53, p27, and p21 expression, we could not detect the expression of p53 by immunoblotting in MSKMRT-31222 tumors (data not shown), although *TP53* was not deleted or mutated based on MSK-IMPACT analysis. Immunohistochemical analysis of MSKMRT-31222 treated mice showed significantly decreased Ki67 staining (Fig. 4D) confirming selinexor’s primary effect on cell proliferation. Notably, the durability of anti-tumor response was dependent upon continued treatment with selinexor since tumor regrowth was observed upon discontinuation of treatment in both models as confirmed by the increase in the percentage of Ki67 positive cells following treatment interruption in the MSKMRT-31222 model (Fig. 1F and 4A,D). Nonetheless, we observe consistent abrogation of tumor growth in MRT models *in vivo* through XPO1 inhibition.

**Figure 4:**
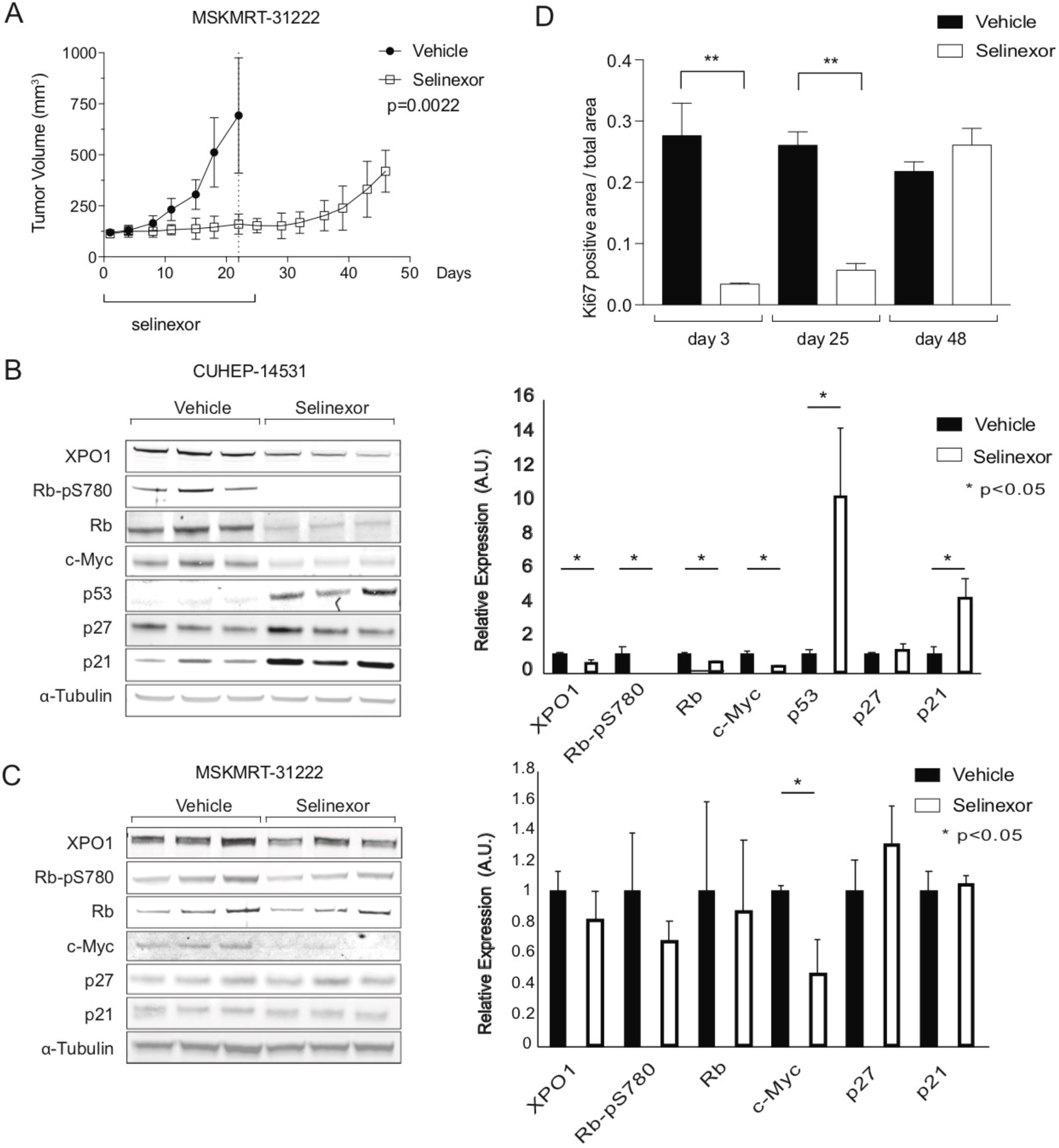
MRT PDX models are sensitive to selinexor *in vivo*. **A)** Tumor response in MSKMRT-31222 PDX (n=8 per arm) treated with selinexor. **B)** CUHEP-14531 and **C)** MSKMRT-31222 immunoblot showing the expression levels of indicated proteins in PDX tumor harvested 3 days following initiation of selinexor or vehicle. Each lane represents tumor from one mouse. Protein expression quantitated by densitometry expressed relative to alpha-tubulin and normalized to vehicle control. (**D)** Immunohistochemistry of PDX tumor following treatment with selinexor evaluating Ki67 staining.

## Discussion

The lack of a standard approach for the treatment of recurrent or progressive disease in MRT and other childhood tumors represents a significant challenge to oncologists. In pediatrics, this challenge is made even more difficult by the paucity of targetable mutations despite increasing access to various next-generation genomic profiling technologies. Hence, novel methodologies are required to interrogate genomic data for non-genetically encoded therapeutic targets. We report on the application of a systems biology approach for identifying non-genetically encoded vulnerabilities in a genomically quiescent cancer within the context of a pediatric precision medicine program. Furthermore, we describe the patient-driven discovery of XPO1 as a novel MRT vulnerability and demonstrate the potential utility of XPO1 inferred activity as a biomarker of response to XPO1 inhibition to identify additional disease indications that would not otherwise be considered for treatment with an XPO1 inhibitor. This experience highlights that although a patient-centric N-of-1 approach begins by focusing on a single patient, the knowledge gained can drive novel discoveries with impact beyond the individual patient.

Partnering an N-of-1 approach with the use of metaVIPER to identify aberrantly activated, druggable proteins offers a novel strategy for precision oncology therapeutics in pediatrics (14). The current approach of identifying and matching targetable mutations with molecularly-targeted drugs (e.g. NCI-MATCH) have resulted in tepid preliminary results (33). Furthermore, while the FDA has approved over 115 targeted antineoplastic drugs over the past two decades, there are only 12 unique genomic alterations linked as companion diagnostics to these drugs (28). MetaVIPER infers the relative activity of over 6,000 proteins, including ∼400 currently druggable proteins. This allows for the identification not only of proteins that are directly activated by a genomic alteration, but also of proteins that become highly activated as the result of a complex interplay of expression, protein-protein interactions, and post-translational modifications.

Using the metaVIPER algorithm, we have identified XPO1 as a highly active protein in MRT and demonstrated that targeting XPO1 through the use of selinexor is efficacious in preclinical models of MRT. We found sub-micromolar IC_50_ values for selinexor across a panel of MRT human cell lines and evidence of cell cycle arrest and apoptosis. *In vivo* treatment of MRT PDX models with selinexor further confirmed a therapeutic response to selinexor, although the extent of the response varied across models.

Selinexor has demonstrated efficacy in patients with refractory hematologic malignancies, as well as those with advanced or metastatic solid tumors (29,30,34). It is currently being evaluated in adult trials for the treatment of a range of cancers including AML, sarcoma, and other solid tumors, and recently garnered FDA accelerated approval for the treatment of multiple myeloma and diffuse large B-cell lymphoma (DLBCL) (28). Notably, our analysis showed that AML, DLBCL, and multiple myeloma have high XPO1 inferred activity, though not as uniformly high as in MRT, supporting the idea that high XPO1 activity may predict response to selinexor. In a multiple myeloma trial of selinexor and dexamethasone, metaVIPER was used with a linear discriminant analysis classifier that identified a four-protein classifier with high predictive performance of patient response to selinexor while using differential gene-expression data did not produce an effective classifier suggesting another potential use of metaVIPER (35). Selinexor is also being studied in a phase 1 pediatric trial in refractory solid tumors and brain tumors (NCT02323880) and is tolerable in combination with fludarabine and cytarabine in a pediatric phase 1 trial of relapsed or refractory acute leukemia (36). This data on the safety and efficacy in children will be helpful for informing future trials in patients with MRT.

Our case illustrates that MRTs are aggressive pediatric malignancies with poor survival rates despite the use of multimodal chemotherapeutic, radiotherapeutic and surgical interventions. Furthermore, intensification of standard therapy approaches has not improved prognosis highlighting the need for more effective therapeutic approaches. Agents targeting downstream effectors of *SMARCB1* loss are currently under clinical investigation for MRTs. The CDK4/6 inhibitor ribociclib previously demonstrated stable disease in several ATRT patients (37) and is currently under investigation in a phase 1 trial in combination with everolimus in pediatric patients with recurrent or refractory ATRT or other malignant brain tumors (NCT03387020). The Aurora-A kinase (AURKA) inhibitor alisertib is currently being studied in a phase 2 trial in pediatric patients with ATRT or MRT (NCT02114229). The EZH2 inhibitor tazemetostat is currently in early phase trials (NCT02601937 and NCT03213665) including the Pediatric MATCH treatment arm for patients with relapsed or refractory *SMARCB1*-deleted tumors, and recently gained FDA approval to treat epithelioid sarcoma (38). It has been previously established that *SMARCB1* mutant MRTs are dependent on the enzymatic activity of EZH2 and that EZH2 inhibition induced tumor regression in pre-clinical models (39). We show that EZH2 inferred activity positively co-segregates with XPO1 activity in MRTs. However, while EZH2 activity was generally increased in MRT, it was not increased to the same extent, and not as universally as XPO1, ranking as the 17^th^ most dysregulated druggable protein across MRTs compared to 3^rd^ for XPO1 (Fig. S2B and Supplementary Table 3). Interestingly, we found that selinexor treatment not only inhibited XPO1 activity, but also significantly decreased the inferred activity of EZH2 (Fig. S3B), suggesting that part of selinexor’s effects may be due to reduced EZH2 activity, in addition to its effects on cell cycle regulators.

Although selinexor can induce significant anti-tumor responses *in vivo*, therapeutic responses were dependent upon sustained drug exposure, suggesting that combination therapies incorporating selinexor with additional agents may be needed to obtain long-term durable responses. Pre-clinical studies of AML show a potential benefit of selinexor in combination with topoisomerase II inhibitors due to increased nuclear retention of topoisomerase II following XPO1 inhibition (40). Furthermore, selinexor treatment was able to restore TOP2A nuclear localization even in cells that were resistant to topoisomerase inhibitor due to aberrant nuclear export and cytoplasmic localization of TOP2A (41). The topoisomerase II inhibitor, etoposide, may therefore be a viable agent for combination with selinexor since it is a principal component of MRT therapy and TOP2A is identified by metaVIPER analysis to be the most activated protein across MRTs in the TARGET cohort. If selinexor demonstrates clinical activity in MRT, a precision medicine approach that couples selinexor with drugs that target proteins uniquely activated in subsets of MRT may also be a viable next step in developing rational combinations that may exert a more durable therapeutic response. In addition, the finding that selinexor treatment increased the activity of particular druggable proteins, including BRAF, PIK3CA, and cereblon, suggests potential targets that can be tested in combination with selinexor to improve response.

In conclusion, we identify XPO1 as a high aberrantly activated protein in MRT and a promising non-genetically encoded therapeutic target. These findings demonstrate the value of metaVIPER analysis in tumors lacking actionable oncogenic alterations to identify potential therapeutic targets and biomarkers of response and illustrate the impact of an N-of-1 approach in identifying viable therapeutic strategies for genomically quiescent tumors beyond the single patient.

XPO1 inhibition as a promising therapeutic strategy for the treatment of MRT will be explored further in the context of a clinical trial of selinexor in pediatric patients with MRT or Wilms tumor, another pediatric tumor demonstrated to have high inferred XPO1 activity.

## Methods

### Cell Culture

Rhabdoid cell lines G401 (ECACC Cat# 87042204, RRID:CVCL_0270) and A204 (ATCC Cat# HTB-82, RRID:CVCL_1058) were obtained from the American Type Culture Collection (ATCC). Rhabdoid cell lines MP-MRT-AN (RRID:CVCL_7049), KP-MRT-NS (RRID:CVCL_7050), KP-MRT-RY (RRID:CVCL_7051), and KP-MRT-YM (RRID:CVCL_7052) were obtained courtesy of Dr. H. Hosoi (42-45). The ATRT cell line CHLA-266 (RRID:CVCL_M149) was obtained from the COG cell line repository. Cell lines were authenticated using STR profiling. Cells were cultured in DMEM with 10% FBS and 1% Anti-Anti (Thermo Fisher Scientific). All cells were tested for *Mycoplasma* approximately once every 4 months (last performed May 2018) (46).

### Cell viability

Cells (2,500 cells/well) were seeded in 96-well plates and after 24 hours were incubated with selinexor (Selleckchem) for 72 hours. The viability of the cells was determined using CellTiter-Glo Luminescent Cell Viability Assay (Promega) according to the manufacturer’s recommendations. IC_50_ concentrations were determined using GraphPad Prism Software Version 7.0.

### Cell cycle assay

Cells were treated with 400 nM selinexor for 48 hours, fixed in 70% ethanol and stored at 4° C for 24 hours. Cells were then washed and stained in 1 × PBS containing 50 µg/ml RNase H (Sigma Aldrich) and 50 µg/ml propidium iodide (PI) (Sigma Aldrich), followed by flow cytometric analysis.

### Apoptosis assay

Cells were treated with 200 nM selinexor for 48 hours. Apoptosis of cells was determined using Caspase-Glo 3/7 Assay (Promega) according to the manufacturer’s recommendations.

### Immunoblotting

Cells and PDX tumor samples were resuspended in 1% SDS-RIPA Buffer (Sigma Aldrich). Tissues were disrupted and homogenized with a Qiagen Bioruptor for 2 minutes at 50Hz (2 rounds). Protein concentrations were determined using the BCA Protein Assay Kit (Pierce) according to manufacturer’s recommendations. The isolated protein samples were separated on 4-12% SDS-PAGE gels and were subsequently transferred to a nitrocellulose membrane using the iBlot Gel Transfer Device and iBlot Gel Transfer Stacks (Thermo Scientific). The blots were developed using LI-COR secondary antibodies (680RD and 800CW). Antibodies are listed in supplementary materials and methods

### Ethics approval and consent to participate

Informed and signed consent was obtained and archived for the research performed and publication of the results. Memorial Sloan Kettering Cancer Center (MSKCC) patients were enrolled onto the MSKCC targeted gene sequencing research study [Genomic profiling in cancer patients; NCT01775072] with approval from the MSKCC Institutional Review Board under protocol IRB# 12-245, #06-107, and #17-387. For xenografts generated at Columbia University Medical Center, the patient was consented for clinical sequencing at Columbia University Medical Center (CUMC) through the Precision in Pediatric Sequencing Program (PIPseq) and PDX tumor mouse model generation under the CUMC Institutional Review Board (IRB)-approved protocol AAAN8404.

### Animal Studies

All experiments were performed in accordance with institutional guidelines and under an approved protocol from the CUMC Institutional Animal Care and Use Committee (Protocol AAAF5850) or the MSKCC Institutional Animal Care and Use Committee (#16-08-011).

### Patient-derived xenograft and xenograft generation

The patient-derived xenograft (PDX) models (CUHEP-14531 and MSKMRT-31222) were developed by transplanting a 2 mm fragment from a patient’s biopsy specimen into the flank of NOD (NOD.Cg-Prkdc^scid^ Il2rg^tm1Wjl^/SzJ) SCID gamma (NSG) mice to generate the passage 0 (P0) generation. Tumors were serially propagated and expanded from the PDX P0 generation for therapeutic studies. The G401 cell line xenograft models were developed by injecting 2 million G401 cells into the flank of NSG mice. Tumor growth was measured biweekly using calipers. Treatment started when flank tumors achieved a tumor volume (TV) of ∼ 150 mm^3^ (TV = width^2^ * length/2). PDX models were dosed with selinexor by oral gavage three times a week at a starting dose of 15 mg/kg which escalated to 20 mg/kg in the second week. Tumors were collected and fragments were either fixed in 10% formalin for histologic analysis or snap frozen in liquid nitrogen for subsequent DNA, RNA, and protein isolation and analyses.

### Immunofluorescence

Immunofluorescence staining was performed in the Molecular Cytology Core Facility of MSKCC using the Discovery XT processor (Ventana Medical Systems). The tissue sections were deparaffinized with EZPrep buffer (Ventana Medical Systems), and antigen retrieval was performed with CC1 buffer (Ventana Medical Systems). Sections were blocked for 30 minutes with Background Buster solution (Innovex), followed by avidin-biotin blocking for 8 minutes (Ventana Medical Systems). Sections were incubated with human-specific anti-Ki67 (DAKO, cat# M7240) or human specific anti-ClCaspase3 (Cell Signaling, cat# 9661), followed by a 60 minute incubation with biotinylated horse anti-mouse IgG (Vector labs, cat# MKB-22258) or biotinylated goat anti-rabbit (Vector Labs, cat# PK6101) respectively. The detection was performed with Streptavidin-HRP D (part of DABMap kit, Ventana Medical Systems), followed by incubation with Tyramide Alexa Fluor 568 (Invitrogen, cat# T20949) prepared according to manufacturer’s instructions with predetermined dilutions. After staining, all slides were counterstained with DAPI (Sigma Aldrich, cat# D9542) for 10 min and cover-slipped with Mowiol (Sigma Aldrich, cat# 81381).

### Nucleic acid extraction, clinical sequencing, and analysis

DNA from frozen tissue or paraffin embedded tissue was extracted using the QIAGEN QIAamp Tissue Kit (for tissue samples) on the QIAcube system. RNA was extracted using the QIAGEN RNeasy Kit (fresh frozen tissue). All slides were evaluated by a pathologist to ensure that a minimum of 50 % viable tumor was present for subsequent extraction and analyses. Whole exome sequencing (WES) was performed using the Agilent SureSelectXT All Exon V5 + UTRs capture kit for library generation and sequenced on the HiSeq 2500 System (Illumina), using paired-end 100 cycle × 2 sequencing. RNA was sequenced using the TruSeq Stranded Total RNA LT Sample Prep Kit (Illumina), with 100 cycles × 2 paired-end sequencing on the HiSeq 2500. DNA sequencing reads were de-multiplexed and converted to FASTQ files using CASAVA from Illumina. Following mapping and variant calling of both tumor and normal samples by NextGENe, resulting variants were subject to filtering. Variants in normal DNA were passed through a “reference range filter” of cancer predisposition genes, genes relevant to pharmacogenomics, and variants relevant to patient care; a “reportable range filter” which includes COSMIC variants in the patient’s mutation report file and variants in genes on the list of ACMG (American College of Medical Genetics and Genomics) recommendations for reporting of secondary findings; as well as a frequency filter. Memorial Sloan Kettering-Integrated Mutation Profiling of Actionable Cancer Targets (MSK-IMPACT) testing was performed as described previously (32).

### MetaVIPER Analysis

All regulatory networks used for metaVIPER analysis were reverse engineered by ARACNe (26). Twenty four core TCGA RNA-Seq derived networks are publicly available in the R Bioconductor package ARACNe networks. In the case of signaling proteins, ARACNe detects maximum information path targets to define its set of “indirectly regulated” target genes.

After standard read alignment of RNA-Seq data by STAR (47) to the GRCh38 reference genome build and summarization of expression quantities at the gene count level, gene expression was normalized by variance stabilization. The stabilization parameters are stored from a single run of the Variance Stabilizing Transformation (VST), as implemented in the DESeq2 package (48), across all TCGA and TARGET samples and these parameters are then applied to individual samples. In the case of novel samples, post-normalization gene expression profiles are compared between the novel samples and tissue-relevant samples from TCGA/TARGET through methods such as principal component analysis and t-SNE to confirm the absence of any significant batch effects that may be attributed to instrument/library-specific biases. In the case of cell line samples, an analogous procedure is performed through the initial application of VST to all CCLE samples.

For the purposes of computing differential protein activity of TARGET MRTs, a gene expression signature (GES) was computed between each MRT versus a pan-cancer reference group that includes gene expression profiles of ∼12,000 tumors belonging to 33 cancer types, followed by application of the analytic Rank-based enrichment analysis (aREA) using each of the available networks (11,12,27). As VIPER is a rank based enrichment method, GES’s are generated through a double-rank transformation process. First, the normalized expression of each gene in the sample is ranked relative to the distribution of the expression of this gene across all TCGA and TARGET samples. Expression distributions for all genes have been fitted to a spline-model to ensure comparable ranks. Next, genes within a sample are ranked based on their relative expressions. Following generation of a GES, aREA tests for a global shift in the positions of a regulatory protein’s target genes in the rank-sorted GES and is an extension of other efforts at computationally efficient method to approximate GSEA (11,49). aREA uses the mean of the rank-transformed positions as the test statistic (enrichment score). The enrichment score is computed first by a 1-tail approach, rank-transforming the absolute values of the GES and secondly by a 2-tail approach, where the positions of the repressed target genes (as inferred in the network) are inverted in the gene expression signature before computing the enrichment score. The contribution of each target gene to the enrichment score is also weighted based on the regulator-target gene mutual information computed by ARACNe. In the final step of aREA, in the case of single sample analysis, statistical significance for the enrichment score is computed by an analytic approach that approximates shuffling the genes in the signature at random, outputting a normalized enrichment score (NES). Gene shuffling is approximated as follows: according to the central limit theorem, the mean of a sufficiently large number of independent random variables is ∼normally distributed. We ensure a mean of zero and variance equal to one for the enrichment score under the null hypothesis by applying a quantile transformation based on the normal distribution to the rank-transformed GES before computing the enrichment score. Then, under the null hypothesis, the enrichment score will be normally distributed with mean of zero and variance *n*^−1^, where *n* is the regulon size, which is further scaled to the standard variable. In the single sample analysis, aREA cannot directly account for biological correlation between the expression of various genes in the given biological context, and so conservative NES/p-value thresholds are empirically used with the assumption that this approach underestimates the p-value. In the final step of metaVIPER analysis, NES’s generated by the application of VIPER using each of the available networks are integrated at the individual regulatory protein level by a weighted average, with greater absolute NES’s contributing a greater weight to the final integrated score to maximize the contribution of signal over noise, as bench marked in (27). As noted, alternative methods including a simple average of NES scores appear to provide an almost equivalent result.

Similarly, metaVIPER was used to infer differential protein activity between samples from each of four selinexor treated MRT/ATRT cell lines and corresponding DMSO treated controls. As these experiments were done in replicate, the GES is computed through gene-wise modified student t-test between condition and control. Pathway analysis on an integrated differential protein activity signature in MRT/ATRT cell lines was performed with ‘Cancer Hallmark’, ‘Gene Ontology’, and ‘Oncogenic Signature’ gene sets provided in the Broad MSigDB collections (50). Pathway enrichment analysis was performed by using the single-tail aREA method described above as a rapid approximation to the KS-test used in GSEA, inputting the pathway genes and the sorted differential protein activity signature. Unlike other statistical tests used for pathway enrichment analysis, such as the Fisher’s exact test on a thresholded list of differentially expressed genes or the repeated application of the Kolmogorov-Smirnov test in classical GSEA, aREA does not require the arbitrary binarization into significant/non-significant hits for target genes in the signature but takes into account their relative position in the signature.

### Co-Segregation Analysis of XPO1 Activity

In order to identify potential genomic events that causally or incidentally associate with increased XPO1 activity, a co-segregation analysis was performed to determine somatic events that are enriched in tumors that are inferred to have high XPO1 activity by metaVIPER across 33 TCGA cancer type cohorts. Publicly available somatic SNV and CNV (GISTIC) calls from the Broad TCGA Genome Data Analysis Center (GDAC) Firehose were downloaded in order to perform this analysis. Enrichment analysis by aREA was performed by arranging tumors in order of lowest to highest XPO1 activity; a null model derived from enrichment analysis of each genomic event with all proteins that metaVIPER infers activity on was developed to estimate the statistical significance of the co-segregation. Analogously, the co-segregation analysis was performed between increased XPO1 activity and the activity of about 411 other ‘druggable’ proteins in both the TARGET malignant rhabdoid tumor dataset and the TCGA datasets to develop hypotheses on biologically relevant protein interactions and potential synergistic drug combinations to treat tumors with elevated XPO1 activity.

## Acknowledgments

This research was supported by NIH grant U01 CA217858 to Andrea Califano (sub-award to Andrew Kung) and NIH grants S10 OD012351 and S10 OD021764 to Andrea Califano.

## Legend

**Supplementary Figure 1:**
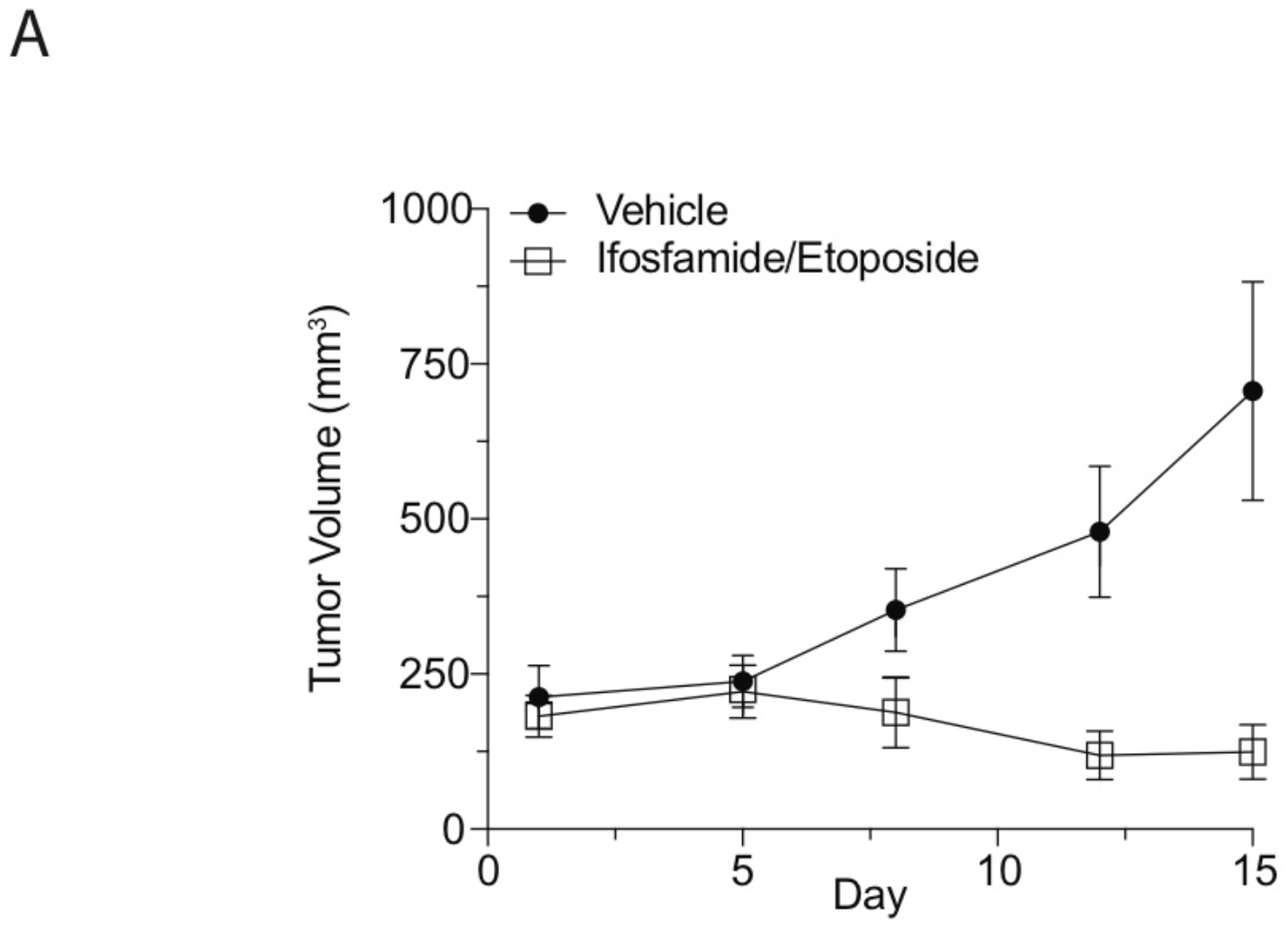
Tumor response of a malignant rhabdoid tumor (MRT) PDX (CUHEP-14531) treated with ifosfamide/etoposide for 15 days.

**Supplementary Figure 2:**
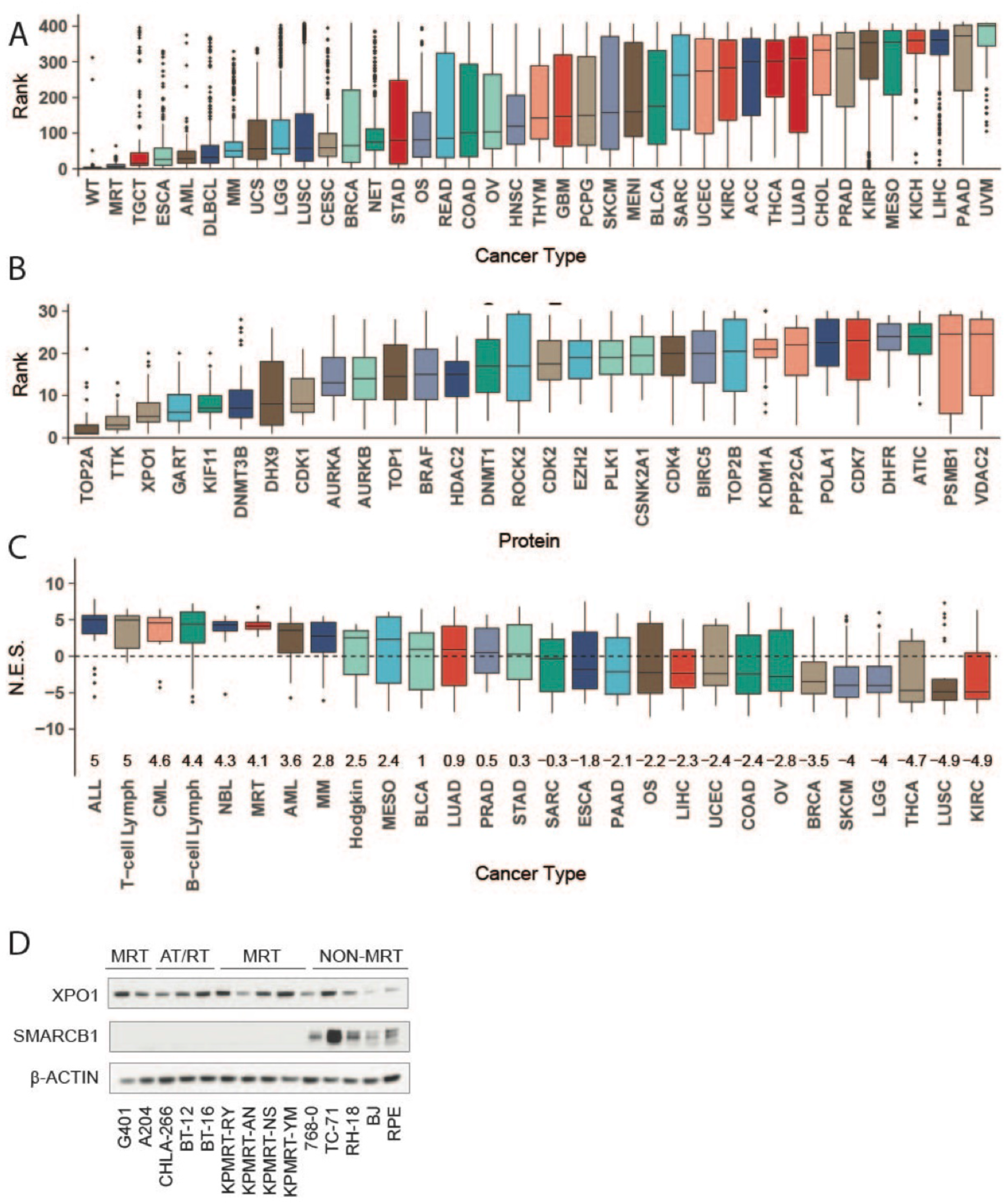
**A)** Boxplots representing the distribution of rank of XPO1 activity relative to 412 ‘druggable’ proteins for which metaVIPER infers activity for 33 TCGA tumor cohorts, 3 TARGET pediatric tumor cohorts (MRT, Wilms, osteosarcoma), and a neuroendocrine tumor cohort. **B)** Boxplots representing inferred activity of the top 30 ‘druggable’ proteins in 68 MRTs in TARGET. **C)** Boxplots representing the distribution of metaVIPER inferred XPO1 activity in a cohort of nine rhabdoid (MRT and ATRT) cell lines compared to 27 cohorts of cancer cell lines profiled in the publicly available CCLE. The median and interquartile range for NES values is represented by each box for the respective cell line cohort. NES values from enrichment analysis are comparable to Z-scores, with higher scores representing increased activity. **D)** Protein expression of XPO1 and SMARCB1 across MRT, ATRT, and non-MRT cell lines.

**Supplementary Figure 3:**
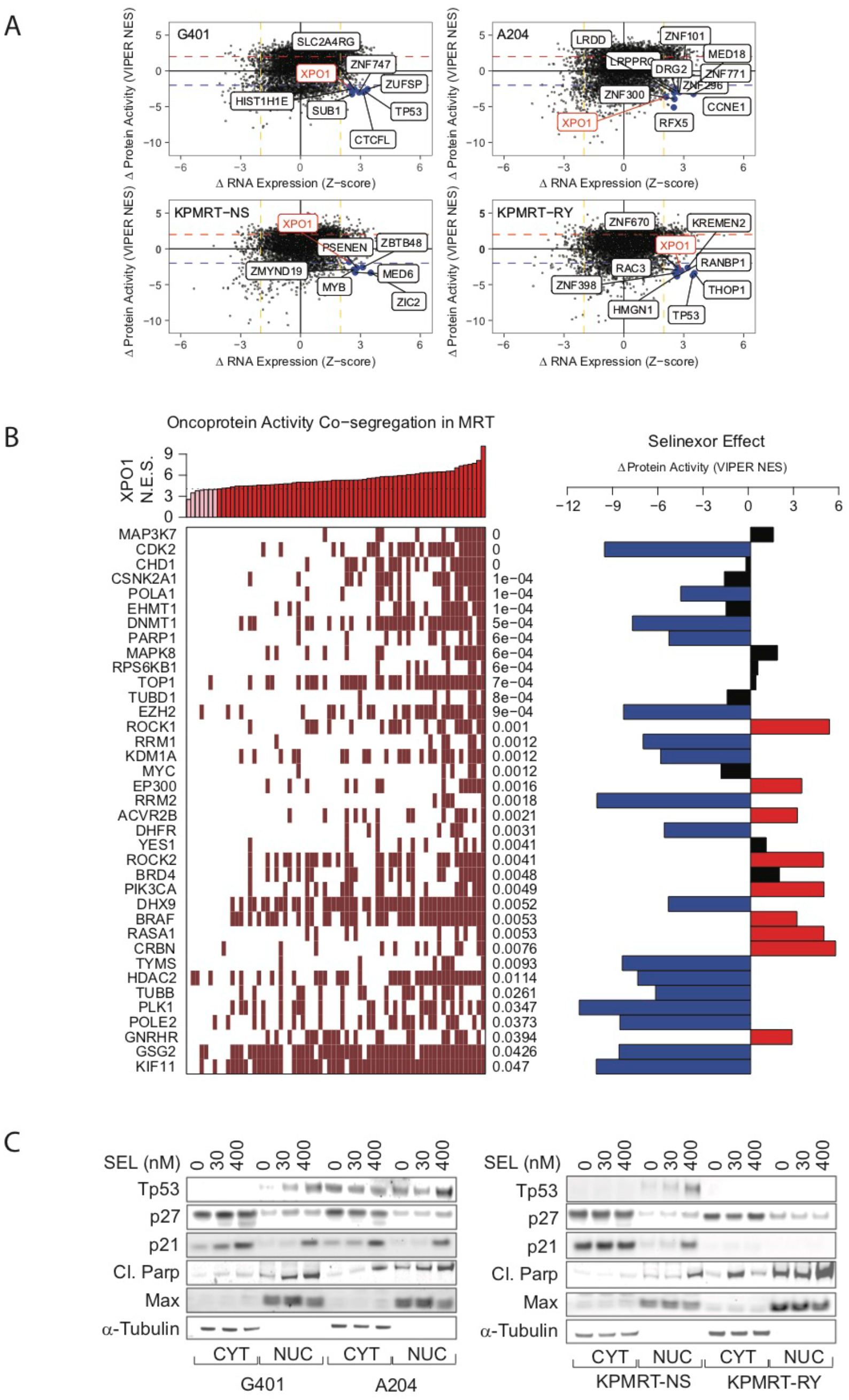
**A)** RNA-Seq was performed on four rhabdoid cell lines at baseline and after treatment with selinexor 30 nM at 6 and 24 hours. Changes in mRNA abundance positively correlate with MetaVIPER inferred activity at 24 hours in each cell line, but XPO1 is amongst a small group of proteins (labeled) whose activity is markedly decreased by selinexor in spite of significant compensatory increase in mRNA. The data represented in Figure 3A is summarized across all four cell lines shown independently here. **B)** Co-segregation analysis on protein activity in 68 rhabdoid tumors from TARGET. Rhabdoid tumors with low to high XPO1 activity are organized from left to right. Dark red boxes represent significant (FDR p <0.01) activation of the respective protein. Selinexor treatment decreases the activity of the majority of proteins whose activity co-segregates with elevated XPO1 activity. **C)** Immunoblot showing expression levels and subcellular localization of the indicated proteins in G401, A204, KPMRT-NS, and KPMRT-RY cells following selinexor treatment.

**Supplementary Table 1**: MetaVIPER inferred activity of 412 ‘druggable’ proteins in the patient tumor biopsy from which the CUHEP-14531 PDX model was developed.

**Supplementary Table 2:** MSK-IMPACT results of MRT cell lines and PDX samples.

**Supplementary Table 3**: MetaVIPER inferred activity of 412 ‘druggable’ proteins in MRT. The mean rank position of XPO1 activity compared to all druggable proteins amongst the 68 rhabdoid tumors in TARGET is provided in the table.

## Notes

Conflict of Interest: Dr. Califano is founder, equity holder, consultant, and director of DarwinHealth Inc., a company that has licensed some of the algorithms used in this manuscript from Columbia University. Columbia University is also an equity holder in DarwinHealth Inc. Dr. Kung is on the Scientific Advisory Board for Karyopharm.

